# Phylogenomic Analyses Support Traditional Relationships within Cnidaria

**DOI:** 10.1101/017632

**Authors:** Felipe Zapata, Freya E. Goetz, Stephen A. Smith, Mark Howison, Stefan Siebert, Samuel H. Church, Steven M Sanders, Cheryl Lewis Ames, Catherine S. McFadden, Scott C. France, Marymegan Daly, Allen G. Collins, Steven H.D. Haddock, Casey W. Dunn, Paulyn Cartwright

**Affiliations:** Department of Ecology and Evolutionary Biology, Brown University, Providence, Rhode Island, United States of America; Computing and Information Services, Brown University, Providence, Rhode Island, United States of America; Department of Ecology and Evolutionary Biology, University of Kansas, Lawrence, Kansas, United States of America; Department of Invertebrate Zoology, Smithsonian Museum of Natural History, Washington District of Columbia, United States of America; Biological Sciences Graduate Program, University of Maryland, College Park, Maryland, United States of America; Department of Biology, Harvey Mudd College, Claremont, California, United States of America; Department of Biology, The University of Louisiana at Lafayette, Lafayette, Louisiana, United States of America; Department of Evolution, Ecology and Organismal Biology, Ohio State University, Columbus, Ohio, United States of America; National Systematics Laboratory of NOAA’s Fisheries Service, National Museum of Natural History, Washington, District of Columbia, United States of America; Monterey Bay Aquarium Research Institute, Moss Landing, California, United States of America

## Abstract

Cnidaria, the sister group to Bilateria, is a highly diverse group of animals in terms of morphology, lifecycles, ecology, and development. How this diversity originated and evolved is not well understood because phylogenetic relationships among major cnidarian lineages are unclear, and recent studies present contrasting phylogenetic hypotheses. Here, we use transcriptome data from 15 newly-sequenced species in combination with 26 publicly available genomes and transcriptomes to assess phylogenetic relationships among major cnidarian lineages. Phylogenetic analyses using different partition schemes and models of molecular evolution, as well as topology tests for alternative phylogenetic relationships, support the monophyly of Medusozoa, Anthozoa, Octocorallia, Hydrozoa, and a clade consisting of Staurozoa, Cubozoa, and Scyphozoa. Support for the monophyly of Hexacorallia is weak due to the equivocal position of Ceriantharia. Taken together, these results further resolve deep cnidarian relationships, largely support traditional phylogenetic views on relationships, and provide a historical framework for studying the evolutionary processes involved in one of the most ancient animal radiations.

## Introduction

Cnidaria is a group of primarily marine invertebrates composed of about 11,000 described species [1] that include reef-forming corals, sea anemones, soft corals, jellyfish, marine hydroids, and freshwater *Hydra* (Fig. 1). Cnidarians are united by the presence of complex intracellular structures called cnidae, with the most universal and diverse cnidae being the stinging structures called nematocysts. The body of cnidarians is, in its simplest form, constructed of two epithelial layers separated by an extracellular mesoglea. Cnidarians are one of the most diverse groups of animals in terms of morphology, lifecycles, ecology, and development. While they are often presented as “simple” animals, many features of presumed simplicity are actually based on misunderstandings of their biology. For example, it is often asserted that cnidarians are radially symmetrical, but most have bilateral symmetry, some have directional asymmetry, and only a subset of species have radial symmetry [2,3]. Fortunately because recent analyses confirm Cnidaria as the sister group to Bilateria [4], the most intensively studied group of animals, we have an excellent outgroup for understanding cnidarian biology.

**Fig. 1.**
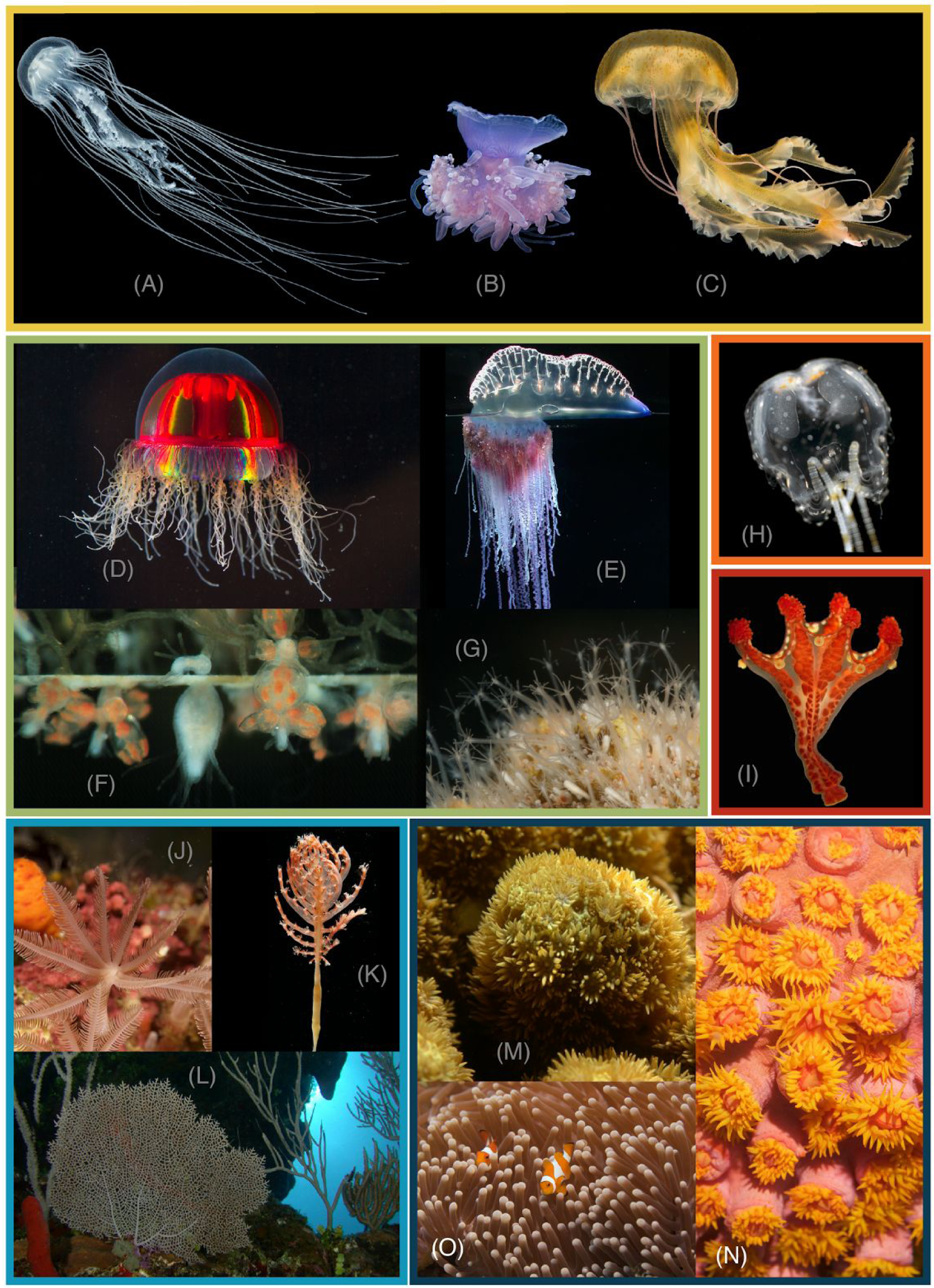
Photographs of cnidarian representatives. The color of the boxes corresponds to the color of clades in Fig. 4 and supplemental figures. (A) Scyphozoa, Pelagiidae: *Chrysaora quinquecirrha*. (B) Scyphozoa, Cepheidae: *Cephea cephea*. (C) Scyphozoa, Pelagiidae: *Pelagia noctiluca*. (D) Hydrozoa, Trachylinae: *Crossota millsae*. (E) Hydrozoa, Siphonophora: *Physalia physalis*. (F) Hydrozoa, Filifera: *Podocoryna carnea*. (G) Hydrozoa, Filifera: *Hydractinia*. (H) Cubozoa: *Copula sivickisi*. (I) Staurozoa: *Haliclystus californiensis*. (J) Octocorallia, Clavulariidae: *Clavularia* sp. (K) Octocorallia, Pennatulidae: *Pennatula* sp. (L) Octocorallia, Gorgoniidae: *Gorgonia ventalina*. (M) Hexacorallia, Poritidae: *Porites* sp. (N) Hexacorallia, Dendrophylliidae: *Tubastrea faulkneri*. (O) Hexacorallia, Stichodactylidae: *Heteractis magnifica*. Photo credits: S. Siebert (A-D), P. Cartwright (F), A. Collins (H-I), and C. Dunn (E, G, J-N).

Cnidaria comprises two groups, Anthozoa and Medusozoa (Fig. 2a). These clades are widely recovered in phylogenetic analyses of molecular data [5–7] (but see [8]) and are supported by morphological characters (e.g., [7,9,10]). Resolving major relationships within Anthozoa and Medusozoa has received considerable attention, but has proven to be challenging (e.g., [11–13]). At least part of that challenge is due to the ancient divergences within Cnidaria. Some fossil representatives from major cnidarian lineages from the Cambrian appear remarkably similar to extant forms [14]. The existence of these crown group Cambrian fossils suggests that multiple extant cnidarian clades already existed over 500 million years ago [15]. The deep and presumably rapid divergence times within Cnidaria, coupled with extensive extinction [16], may present a particularly difficult hurdle when reconstructing higher level phylogenetic relationships within this group.

**Fig. 2.**
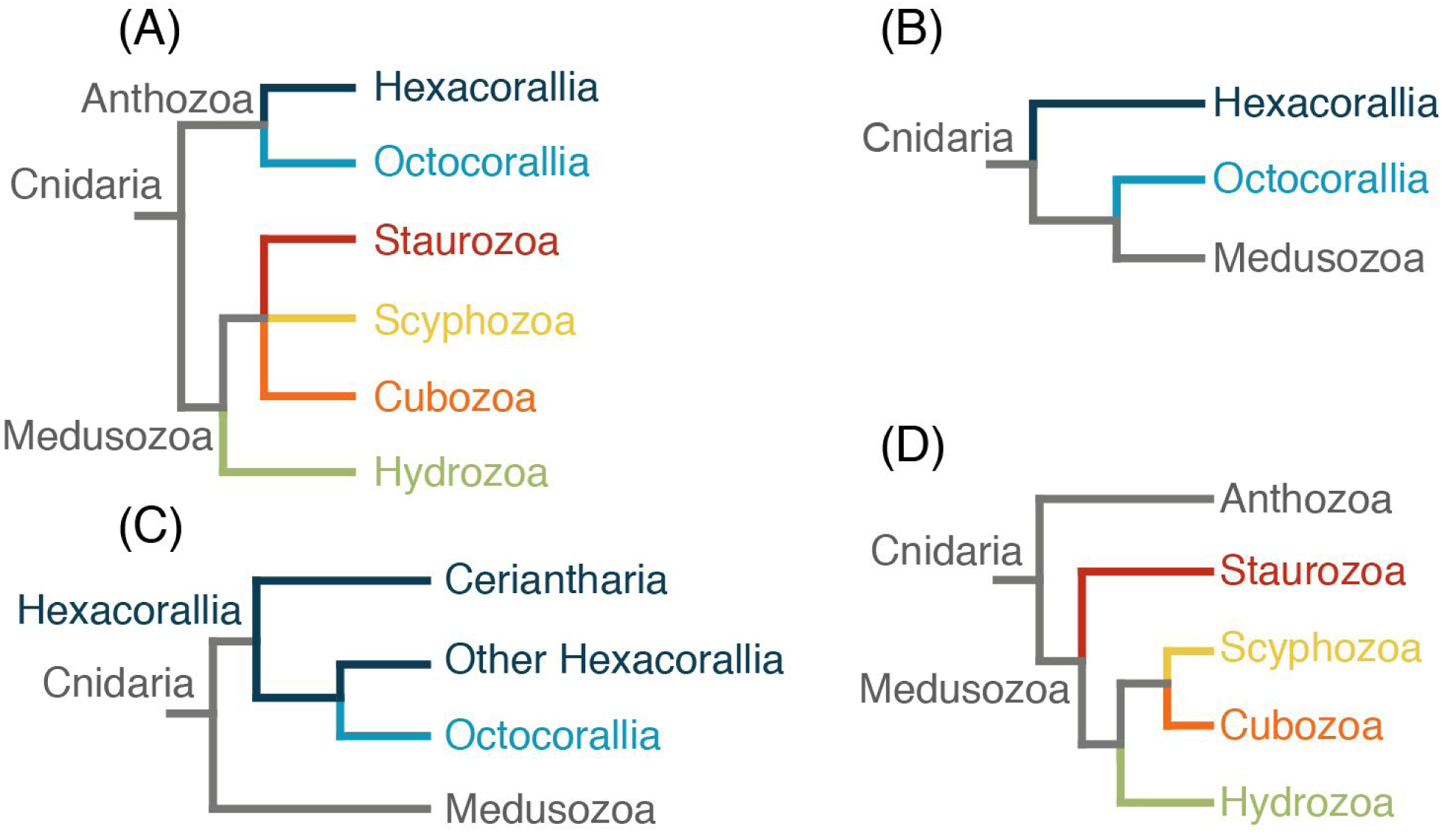
Alternative hypotheses for internal relationships within Cnidaria. (A) Traditional classification and relationships within Cnidaria. (B) Anthozoa paraphyletic with Octocorallia sister to Medusozoa [8]. (C) Hexacorallia paraphyletic with Ceriantharia sister to Hexacorallia + Octocorallia clade [17]. (D) Staurozoa as the sister taxon to the rest of Medusozoa [7]. The color of the branches corresponds to the color of clades in Fig. 4 and supplemental figures.

Anthozoa contains approximately 7,500 extant described species [18]. It is composed of two major groups, Hexacorallia (sea anemones, tube anemones, scleractinian corals, and black corals) and Octocorallia (soft corals, gorgonians, and sea pens). Studies using nuclear ribosomal DNA markers recover anthozoan monophyly [6,17,19–21]. Morphological synapomorphies that support anthozoan monophyly include the actinopharynx, which is an ectodermally-lined tube that extends from the mouth into the gastric cavity, the siphonoglyph, which is a ciliated groove in the actinopharynx, and the mesenteries, which are sheets of gastrodermal tissue that extend from the body wall into the coelenteron and have musculature of gastrodermal origin [18,22,23]. All anthozoans also have bilaterally symmetric polyps [24,25]. Whether any of these morphological features are plesiomorphic for Cnidaria cannot be distinguished in the absence of a robust phylogeny. This issue is confounded by recent molecular phylogenetic studies using mitochondrial genome sequences that recover a paraphyletic Anthozoa, with Octocorallia as the sister taxon to Medusozoa [8,26] (Fig. 2b).

Resolving deep relationships within Anthozoa has been difficult. Octocoral polyps have eight tentacles, eight mesenteries, and almost all species are colonial. They also have a unique gene, *mtMutS,* in their mitochondrial genome [27,28]. Several molecular studies support the monophyly of Octocorallia [19,21,22,29,30]. Although Octocorallia is traditionally divided into three groups, Pennatulacea (sea pens), Helioporacea (blue corals) and Alcyonacea (soft corals and gorgonians), Alcyonacea is likely paraphyletic, as are many of the traditionally defined groups within it [11,31]. Molecular phylogenetic studies of Octocorallia have converged on three well-supported lineages, all of which include representatives from Alcyonacea (reviewed in [31]): the Holaxonia-Alcyoniina group, the Calcaxonia-Pennatulacea group, which includes a paraphyletic Calcaxonia, with a monophyletic Pennatulacea and Helioporacea, and the *Anthomastus*-*Corallium* clade, which includes representatives from Scleraxonia and Alcyoniina. Hexacorals are diverse in polyp morphology and organization, including colonial and solitary species that have bodies with tentacles and mesenteries in multiples of six, eight, or ten. All hexacorals possess a distinct type of stinging organelle (cnida) called a spirocyst [32]. In contrast to octocorals, the traditional ordinal groups of hexacorals are monophyletic [18]. Hexacoral monophyly has been supported by several molecular studies [22,25,30]. The molecular phylogenies in which hexacorals are monophyletic all recover the tube anemones (order Ceriantharia) as sister to the rest of hexacorals. However, this finding has been challenged recently by Stampar *et al.* [17], who found Ceriantharia as sister to all other anthozoans, rendering Hexacorallia paraphyletic (Fig. 2c).

Medusozoa comprises approximately 3,700 extant described species and is usually divided into four groups, Scyphozoa (true jellyfish), Cubozoa (box jellies), Staurozoa (stalked jellyfish), and Hydrozoa (hydroids, hydromedusae, siphonophores) [18]. While medusozoans are often thought of as being characterized by the presence of a free-swimming medusa stage, this is far from universal within the group [7,31]. Instead, all medusozoans have a linear mitochondrial DNA genome [5,10] and a hinged cap called an operculum at the apex of their nematocysts [23]. These synapomorphies are consistent with the monophyly of Medusozoa recovered by molecular phylogenetic studies using nuclear ribosomal DNA sequences [7,15,33]. Symmetry is quite diverse in Medusozoa. Different species display bilateral or radial symmetry, and some even exhibit directional asymmetry [2,3,34].

Relationships among major medusozoan lineages have received inconsistent support, and some findings remain controversial. These include the rooting of Medusozoa with regard to the position of Staurozoa [7,33], and the sister relationship between Scyphozoa and Cubozoa. Staurozoa comprises about 50 species that have long been confusing to cnidarian systematists due to their benthic polyp forms that also exhibit characters known in the medusa stages of cubozoans and scyphozoans, such as gastric filaments, coronal muscle, and structures derived from primary tentacles of the polyp (rhopalioids/rhopalia). Maximum-likelihood analyses of nuclear ribosomal sequences recover Staurozoa as the sister taxon to the rest of Medusozoa, and a monophyletic Cubozoa and Scyphozoa group as sister to Hydrozoa [7,15] (Fig. 2d). These results are contradicted by an analysis of protein coding mitochondrial gene sequences, which recovered a paraphyletic Scyphozoa and a Staurozoa and Cubozoa clade as the sister taxon to Hydrozoa [8]. In a cladistic analysis of morphological data, Marques and Collins [9] report Cubozoa and Staurozoa as sister to Scyphozoa, whereas an analysis of a corrected version of the same dataset was consistent with the results derived from nuclear ribosomal sequences [35]. Resolving the relationships among these lineages has implications for our understanding of key innovations within Medusozoa, including the origin of a pelagic medusa and associated sensory structures and swimming musculature, as well as mode of medusae metamorphosis and development.

Here, we present a broadly sampled phylogenomic analysis of Cnidaria designed to test the general framework for cnidarian phylogeny that has emerged in the past decades, and compare alternative hypotheses for remaining questions. By collecting new transcriptome data for 15 species and analysing them in conjunction with publicly available transcriptomes and genomes, we present a robust hypothesis of higher-level relationships in Cnidaria.

## Materials and Methods

### Taxon sampling, RNA isolation, and Sequencing

New transcriptome data were sequenced for 15 species using Roche 454 GS FLX Titanium and Illumina HiSeq 2000/2500 sequencers. Sample preparation protocol and sequencing technology for each sample are listed in S1 Table. All new data were deposited in the NCBI Sequence Read Archive (BioProject PRJNA263637). In combination with publicly available data, sequences from 41 taxa were used for matrix construction.

### Data analyses

All 454 data were assembled with Newbler (version 2.5.3). Agalma (versions 0.4.0-0.5.0) [36] was used for all other analysis steps through supermatrix construction. A git repository of the analysis code is available at https://bitbucket.org/caseywdunn/cnidaria2014. This source code is sufficient to reconstruct the supermatrix from the data, and includes all settings and parameters used for these intermediate steps. Agalma is a workflow that automatizes all the steps in a phylogenomic analysis, and keeps track of data provenance and parameters used in the analysis, allowing full reproducibility of the results. It takes Illumina sequence reads and after filtering and quality control, it generates fully annotated assemblies. Externally assembled transcriptomes can also be imported into Agalma for downstream analysis. Across species, Agalma identifies homologous sequences, determines gene orthology based on gene tree topology, and generates a supermatrix of concatenated orthologous genes.

We sampled 1,262 genes to generate a supermatrix with 50% occupancy. This matrix has a length of 365,159 aa (Fig. 3). Three taxa, *Calibelemnon francei*, *Craspedacusta sowerbii*, and *Obelia longissima*, had less than 5% occupancy and were excluded from further analyses. The primary matrix (matrix 1) used for all phylogenetic analyses therefore has 38 taxa and 54% gene occupancy. From this matrix, we constructed a reduced matrix (matrix 2) from which two poorly sampled taxa, the ceriantharian (16.6% gene sampling) and *Haliclystus sanjuanensis* (6.5% gene sampling), were also removed since they were unstable in the primary analyses. This produced a reduced matrix with 57% gene occupancy.

**Fig. 3.**
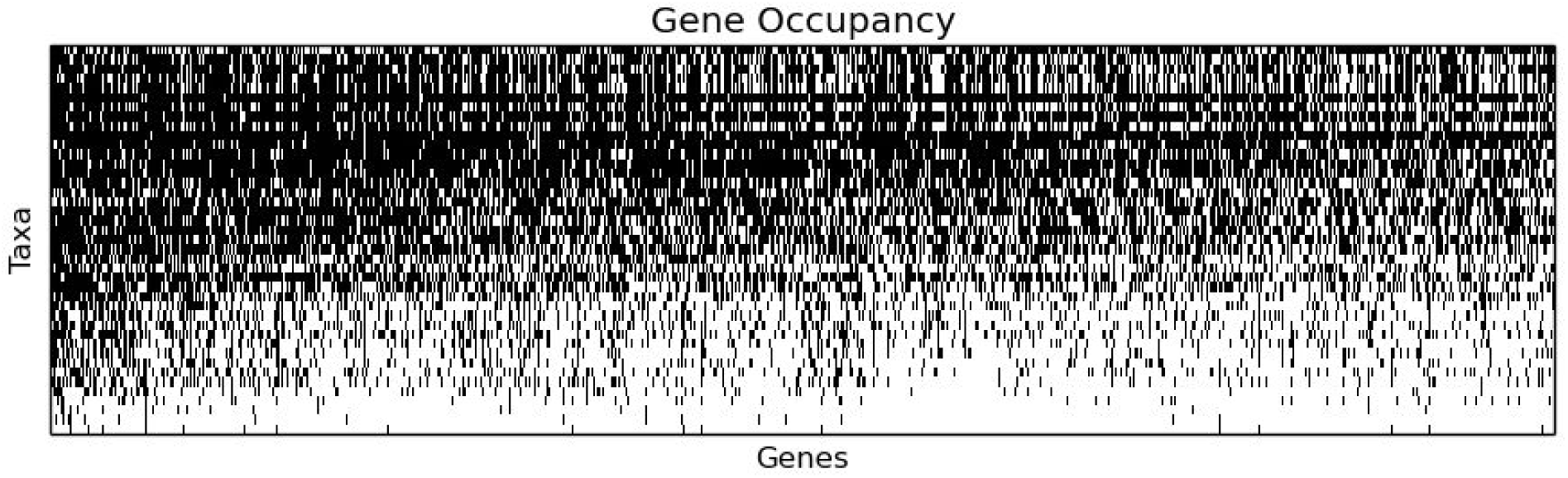
The 50% gene occupancy matrix. Black indicates sampled genes for each of the 41 taxa. Genes and species are sorted by sampling, with the best sampled in the upper left. The last three taxa, *Calibelemnon francei, Craspedacusta sowerbii*, and *Obelia longissima*, had less than 5% gene occupancy and were excluded from further analyses to produce matrix 1.

We inferred phylogenetic relationships using both Maximum Likelihood (ML) and Bayesian Inference (BI) approaches. For ML, we used ExaML v 1.0.12 [37] with the WAG+Γ model of amino acid substitution on the unpartitioned matrices 1 and 2. We also ran a partitioned ML analysis on matrix 1 according to results of PartitionFinder v 1.1.1 [38]. For PartitionFinder, we used genes as initial partitions, linked branch lengths across partitions, used the Bayesian Information Criterion (BIC) to select among all models of amino acid substitution, and used the relaxed hierarchical clustering algorithm to search for a good partitioning scheme. Bootstrap values were estimated on the unpartitioned and partitioned analyses with 200 replicates. BI was conducted on PhyloBayes MPI v. 1.4e [39] using the CAT model of evolution [40] with the global exchange rates fixed to uniform values (CAT-Poisson) and inferred from the data (CAT-GTR). For these analyses, constant sites were removed from the alignment to improve MCMC mixing [39]. Two independent MCMC chains were run on matrix 1, adjusting the number of cycles until convergence was achieved. Analyses with the CAT-GTR setting did not converge despite long CPU time, thus we do not include results from these analyses here. Convergence was determined with time-series plots of the likelihood scores, and maximum bipartition discrepancies across chains less than 0.1. Post-burn-in (50%) sampled trees were combined and summarized with a majority rule consensus tree.

### Hypothesis testing

We used the SOWH test [41] to evaluate three phylogenetic hypotheses: (i) Octocorallia is sister to Medusozoa (i.e., Anthozoa is paraphyletic) [8], (ii) Ceriantharia is sister to the Hexacorallia and Octocorallia clade (i.e., Hexacorallia paraphyletic) [19], and (iii) Staurozoa is sister to all other Medusozoa [33]. To carry out these analyses, we used SOWHAT [42] specifying a constraint tree and the WAG+Γ model on matrix 1. We used the stopping criterion implemented in SOWHAT to determine an appropriate sample size for the null distribution. The commit version at the time we ran these analyses is available at https://github.com/josephryan/sowhat/commit/e0c214e8d7756211d7cbb4a414642c257df6b411

## Results and Discussion

Phylogenetic results are congruent across inference methods, models of molecular evolution, and partitioning schemes (Fig. 4, S1 and S2 Figs). All our analyses provide strong support for the reciprocal monophyly of Anthozoa and Medusozoa, with the placement of the root for Cnidaria between these two clades. The Anthozoa/Medusozoa split is consistent with previous molecular phylogenetic studies based on rDNA sequences [6,7,10] and morphological synapomorphies [9,18]. This result is not consistent with the results of Park *et al.* [26] and Kayal *et al.* [8], which recover Anthozoa as paraphyletic using mitochondrial DNA sequences. A tree enforcing Octocorallia as sister to Medusozoa, rendering Anthozoa paraphyletic (Fig. 2b), is significantly worse (SOWH test: *n* = 100, Δ-likelihood = 2523.533, *p* = 0) than our most likely tree (Fig. 4). This is consistent with Kayal *et al.* [8] who could not reject anthozoan monophyly using any statistical test of topology. If Anthozoa is non-monophyletic, then those features unique to Anthozoa, including the actinopharynx, siphonoglyph, and mesenteries with musculature of gastrodermal origin, would be interpreted as either convergent in Octocorallia and Hexacorallia, or as ancestral features of Cnidaria lost or transformed in Medusozoa. Our results contradict this view and confirm that these features are synapomorphies of Anthozoa. Our results do not recover Coelenterata, a clade comprised of Cnidaria and Ctenophora that has been recovered in some analyses [43]. Removing the ctenophore from the analysis did not alter relationships between other taxa (S4 Fig).

**Fig. 4.**
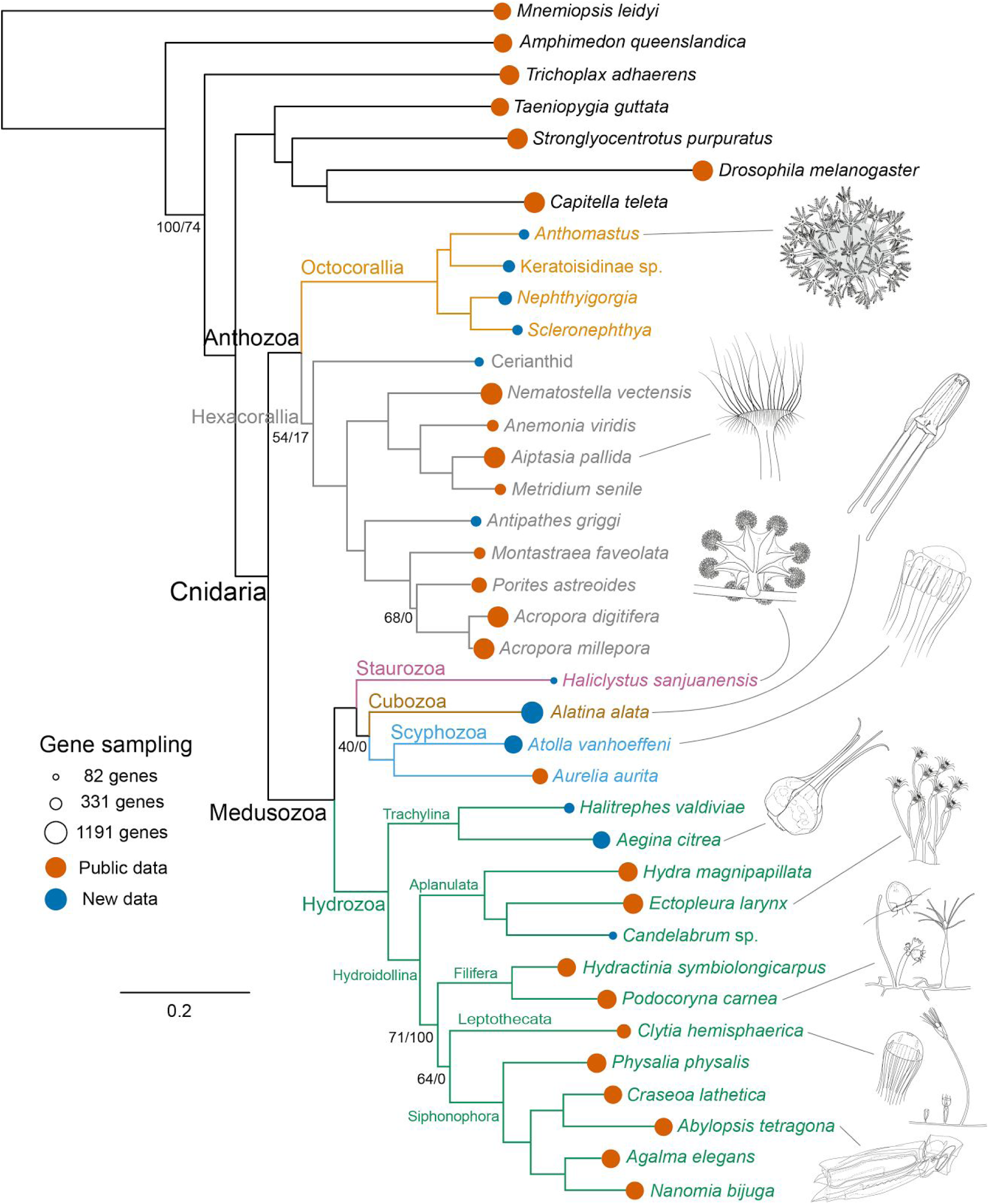
Rooted phylogram of the maximum likelihood (ML) analysis. Branch support values correspond to percent ML-bootstrap values/percent Bayesian posterior probabilities. No values are shown for branches with 100/100 support. The areas of the lollipops, at the branch tips, are proportional to the number of genes sampled. Illustrations (by F. Goetz) are provided for select species, as indicated by lead lines.

Within Anthozoa, the monophyly of Hexacorallia has low support due to the phylogenetic instability of Ceriantharia (Fig. 4, S1 and S2 Figs.), our most poorly sampled taxon within Anthozoa (16.6% gene sampling). Each analysis provides mixed support for the placement of Ceriantharia as either sister to the rest of the Hexacorallia, i.e., Hexacorallia is monophyletic (54% ML, 17% Bayes; Fig. 4, S2 Fig.), or sister to Octocorallia, i.e., Hexacorallia is paraphyletic (46% ML, 81% Bayes; S1 Fig.). Removing Ceriantharia clearly shows the monophyly of all other sampled Hexacorallia (S3 Fig.). The traditional view of hexacoral monophyly (Fig. 4, S2 Fig.) is also supported by previous molecular phylogenetic studies [6,8,25] and compelling morphological synapomorphies (discussed above). In particular, ceriantharians share with hexacorals a unique type of cnida called a spirocyst [32]. A spirocyst is ontogenetically and chemically similar to a nematocyst, and is inferred to have a common origin (see [23]), but it is a single walled capsule whose internal tubule is sticky. No instances of evolutionary losses of cnidae, nematocysts included, have been reported. Stampar *et al.,* [17] also recovered a sister relationship between Ceriantharia and Octocorallia with low support considering only 28S rDNA sequences. However, due to overall better support values, Stampar *et al.,* [17] prefer the topology recovered with 16S rDNA sequences, where Ceriantharia is sister to the rest of the Anthozoa. Enforcing this topology (Fig. 2c) is significantly worse (SOWH test: *n* = 86, Δ-likelihood = 78.0694, *p* = 0) than our most likely tree (Fig. 4). Although not discussed by Stampar *et al.* [17], their interpretation of anthozoan phylogeny requires that spirocysts are lost in Octocorallia. The cnidome of Octocorallia includes only a limited suite of nematocysts (none of which are unique to the group: see [32,44]) and no single-walled cnidae, so it is improbable that these have been transformed into another type of cnida. The alternative explanation for this feature under the preferred phylogeny of Stampar *et al.* [17] is that the spirocysts of Ceriantharia and of other Hexacorallia are convergent.

The monophyly of Octocorallia is strongly supported in all our analyses (Fig.4, S1-S3 Figs.). Although our sampling of octocorals is limited to four taxa, it represents the breadth of our current understanding of octocoral phylogenetic diversity [31]. Specifically, all three major clades of octocorals are represented. These are the Holaxonia - Alcyoniina clade (represented here by *Scleronephthya* and *Nephthyigorgia*), the *Anthomastus - Corallium* clade (represented by *Anthomastus*), and the Calcaxonia Pennatulacea clade (represented by Keratoisidinae sp.). Relationships among these four taxa are congruent with recent octocoral phylogenies [11,31]. Resolution within these deep nodes suggests that this phylogenomic approach should prove valuable to reconstructing higher level octocoral phylogeny as more taxa are analyzed in future studies.

Medusozoa, comprising Scyphozoa, Staurozoa, Cubozoa, and Hydrozoa, forms a strongly supported monophyletic group (Fig. 4, S1-S3 Figs.). All our analyses support a sister group relationship between Hydrozoa and a clade composed of Scyphozoa, Staurozoa, and Cubozoa. This clade revives the traditional sense of Scyphozoa, prior to the elevation of Stauromedusae and Cubomedusae to distinct classes [9,45]. The only staurozoan included in our analysis, *Haliclystus sanjuanensis* (6.5% gene sampling), is the most poorly sampled taxon in our data set (Fig. 4). While all analyses place it within this clade with strong support, its position within the clade is unstable and it moves between positions as sister to Cubozoa and Scyphozoa (40% ML, 0% Bayes; Fig. 4) and sister to Cubozoa (60% ML, 100% Bayes; S1 Fig.). When the staurozoan is excluded from the analyses, the cubozoan *Alatina alata* is sister to the scyphozoans with 100% support (S3 Fig.). Collins *et al.* [7] reported Staurozoa as sister to the rest of Medusozoa, suggesting that pelagic medusae evolved after the divergence of staurozoans. Our results do not support this hypothesis and resulting scenario of medusa evolution. Enforcing the staurozoan as sister to all other medusozoans [33] (Fig. 2d) is significantly worse (SOWH test: *n* = 100, Δ-likelihood = 118.6461, *p* = 0) than our most likely tree (Fig. 4). Instead, our results are consistent with the cladistic analysis of Marques and Collins [9] based on morphology and life history features. Characters from Marques and Collins [9] that support the clade composed of Staurozoa, Cubozoa, and Scyphozoa include radial tetramerous symmetry of the polyp stage, medusa production involving metamorphosis of the oral end of the polyp, canal systems in the polyps, musculature organized in bundles of ectodermal origin, rhopalia or rhopalia-like structures, and gastric filaments. Characters supporting a Cubozoa + Staurozoa clade include quadrate cross section and metamorphosis of medusae without fission [9].

Recovered relationships within Hydrozoa are largely consistent with those found in previous studies [7,12], including the reciprocally monophyletic Trachylina and Hydroidolina. Trachylina is composed of Narcomedusae (represented here by *Aegina citrea*), Trachymedusae (represented here by *Halitrephes valdiviae*), and Limnomedusae (not represented). Within Hydroidolina, our sampling includes representatives of Siphonophora, Aplanulata, “Filifera” (which has previously been shown to be polyphyletic [12,46]), and Leptothecata. Relationships among the major lineages of Hydroidolina have been difficult to resolve [12,46]. The analyses presented here recovered the Aplanulata clade as sister to the rest of the sampled representatives of Hydroidolina. Given that members of Trachylina and Aplanulata are mostly solitary species (see [47]), these results may imply that coloniality in Hydrozoa evolved following the divergence of Aplanulata from the rest of Hydroidolina, as opposed to at the base of Hydroidolina as reported by Cartwright and Nawrocki [46]. It should be noted however that representatives of other colonial hydroidolinan lineages including Capitata and other Filifera were not included in this analysis, so the precise origin of coloniality within Hydrozoa awaits further sampling. The monophyly of Aplanulata and Siphonophora are strongly supported. The internal relationships of Siphonophora are in accord with previously published results [48], while those of Aplanulata differ from previous results [49] in that *Ectopleura* is more closely related to *Candelabrum* than to *Hydra.*

## Conclusions

Although divergences within major lineages of Cnidaria likely occurred over half a billion years ago [14,15], using a phylogenomic approach this study reveals strong support for many deep nodes within the cnidarian tree of life (S5 Fig.). This represents a significant improvement from previous studies using rDNA markers which, in many cases, failed to resolve relationships among major cnidarian clades. Our study is also consistent with more traditional hypotheses of cnidarian relationships including the monophyly of Hexacorallia, Anthozoa, and a clade composed of Staurozoa, Cubozoa, and Scyphozoa. Future phylogenetic studies with increased taxonomic sampling will continue to resolve more detailed relationships and patterns of character evolution in this highly diverse group.

## Acknowledgements

Thanks to Caitlin Feehery for assistance with preparation of some samples. Thanks to Rob Steele for the *Craspedacusta* sample (from a line established by Terry Peard), Tony Montgomery for the *Antipathes* sample, and Claudia Mills for the *Haliclystus* sample. Rebecca R. Helm provided helpful comments on the manuscript.

## Author contributions

PC and CWD conceived of and designed the study. FZ designed and ran analyses. FG extracted and prepared most samples for sequencing, along with SS. SAS assembled the 454 data and performed preliminary analyses. SHC performed the SOWH tests. SMS and PC generated the *Hydractinia* and *Podocoryna* data. CSM and SCF led octocoral sampling, and MD guided hexacoral sampling. CLA, AGC and PC generated the *Alatina alata* data. CM and SF provided the octocorals. MD collected most hexacorals. SHDH collected many of the medusozoans. MH assisted with data management, submission, and analysis implementation. PC, FZ, and CWD wrote the manuscript with considerable input from other authors. FG drew line illustrations of animals. All authors discussed and/or contributed to the final manuscript version.

## Supporting Information

**S1 Fig.**
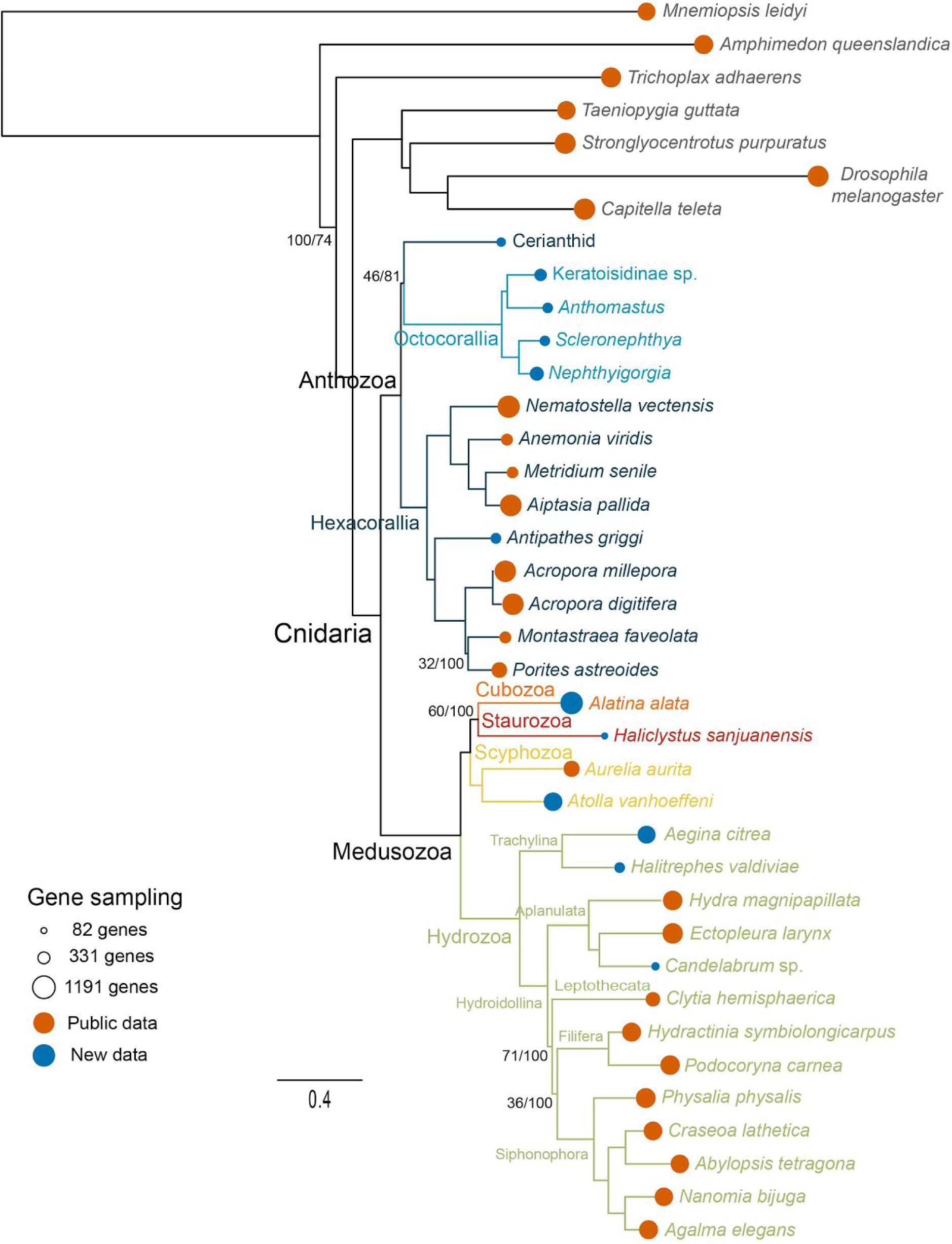
Majority rule consensus rooted phylogram of Bayesian Inference (BI) analysis. Branch support values correspond to percent ML-bootstrap values/percent Bayesian posterior probabilities. No values are shown for branches with 100/100 support. The areas of the lollipops, at the branch tips, are proportional to the number of genes sampled.

**S2 Fig.**
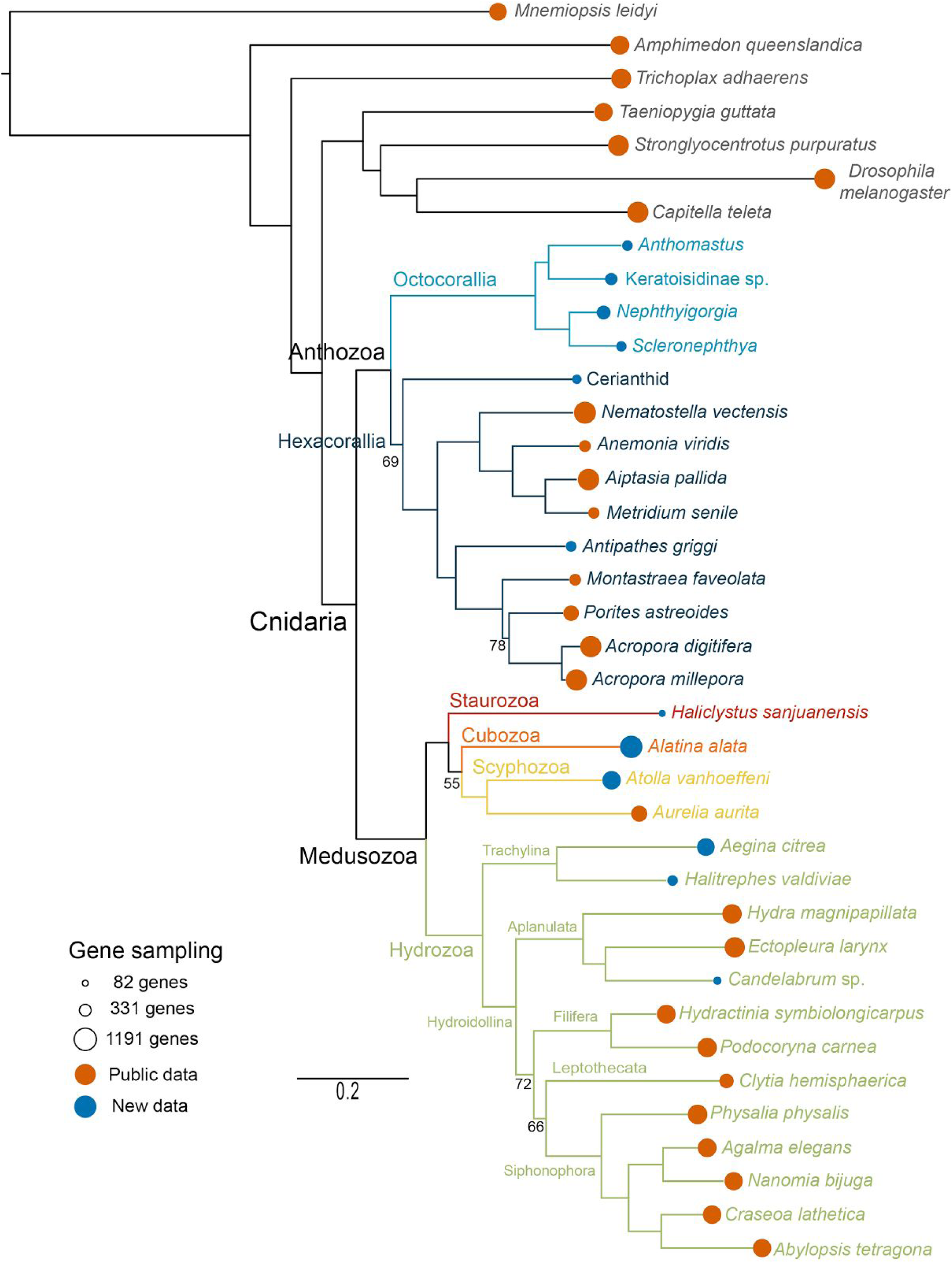
Rooted phylogram of maximum likelihood (ML) partitioned analysis. Branch support values correspond percent bootstraps. No values are shown for branches with 100% support. The areas of the lollipops, at the branch tips, are proportional to the number of genes sampled.

**S3 Fig. 3.**
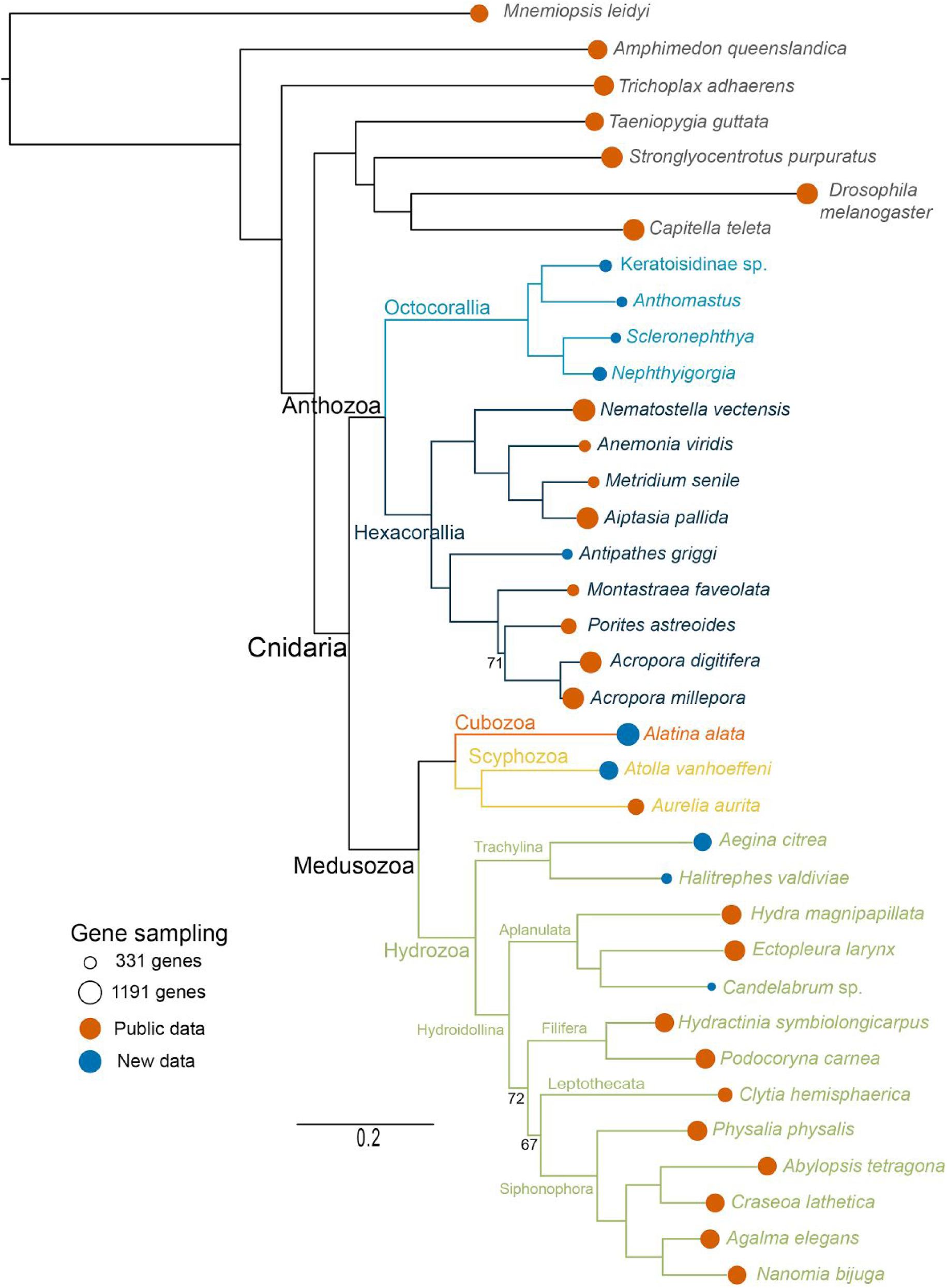
Rooted phylogram of the maximum likelihood (ML) analysis with unstable poorly sampled taxa (*Haliclystus sanjuanensis* and ceriantharian) removed. Branch support values correspond to percent bootstraps. No values are shown for branches with 100% support. The areas of the lollipops, at the branch tips, are proportional to the number of genes sampled.

**S4 Fig.**
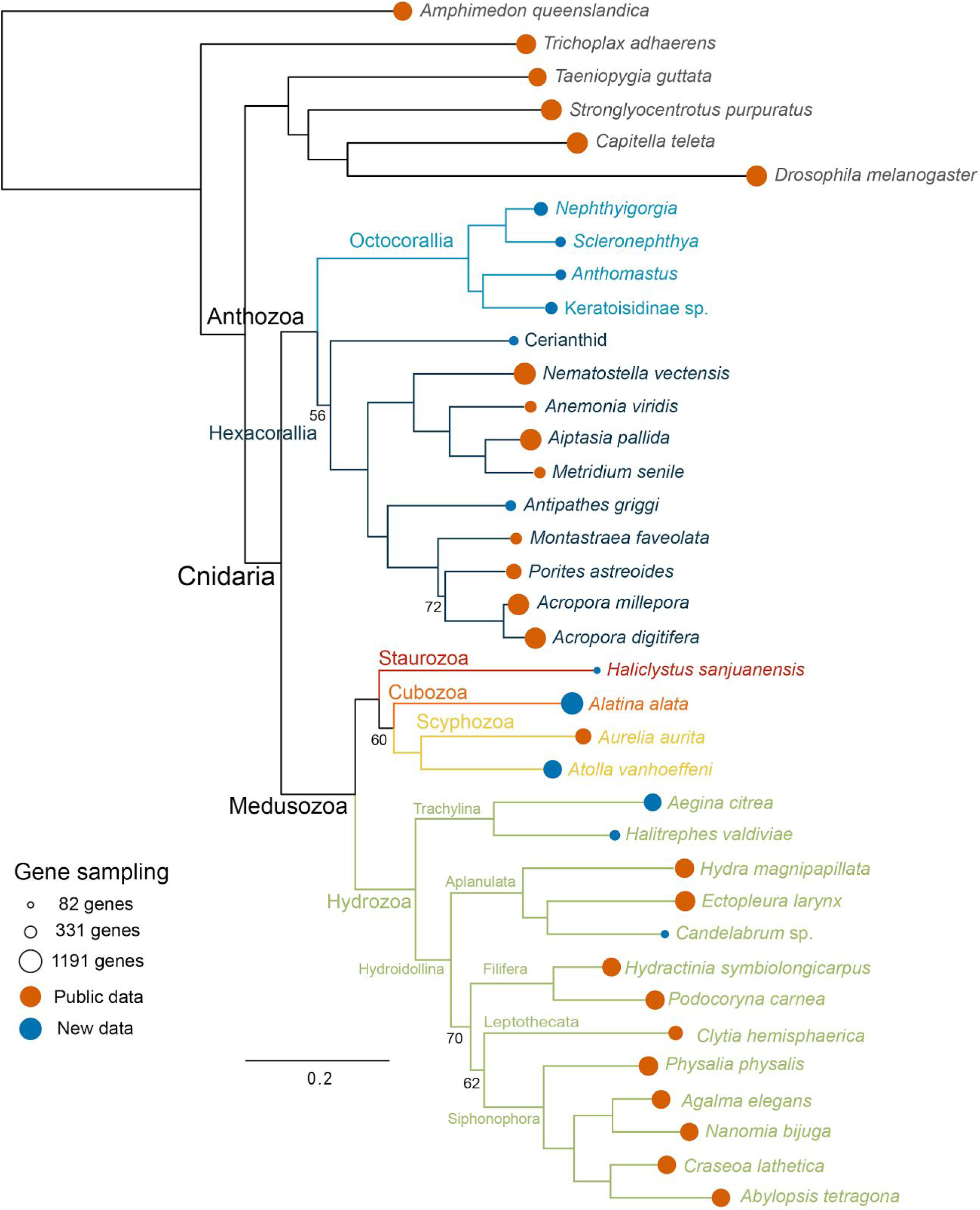
Rooted phylogram of maximum likelihood (ML) with Ctenophore removed. Branch support values correspond percent bootstraps. No values are shown for branches with 100% support. The areas of the lollipops, at the branch tips, are proportional to the number of genes sampled.

**S5 Fig.**
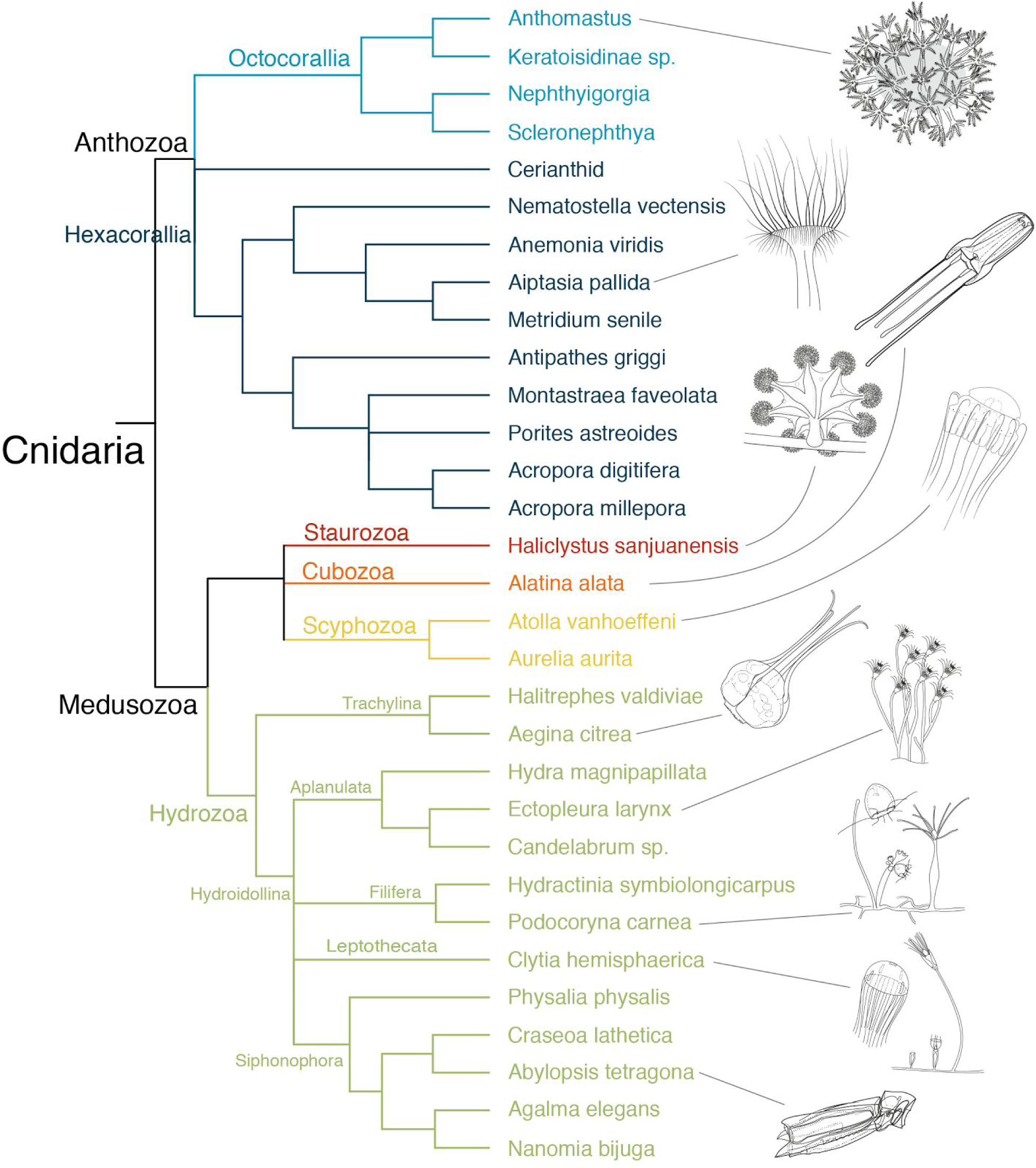
Summary of major findings. Cladogram of Cnidaria based on phylogeny in Fig. 4. Branches that did not receive 100% support in ML and Bayesian analyses are collapsed to polytomies.

**S1 Table. Specimen data.** Accession numbers or URLs for all data considered in this analysis, including data that were previously public and those that are newly generated here. A csv version of this table is available in the git repository (see Data Availability Statement).

## References

1. Appeltans W, Ahyong ST, Anderson G, Angel MV, Artois T, Bailly N, et al. The magnitude of global marine species diversity. Curr Biol. 2012;22: 2189–2202. doi:10.1016/j.cub.2012.09.036

2. Hyman LH. The Invertebrates: Protozoa through Ctenophora. McGraw-Hil Book Company. Inc New York. 1940;

3. Manuel M. Early evolution of symmetry and polarity in metazoan body plans. C R Biol. 2009;332: 184–209. doi:10.1016/j.crvi.2008.07.009

4. Dunn CW, Giribet G, Edgecombe GD, Hejnol A. Animal Phylogeny and Its Evolutionary Implications. Annu Rev Ecol Evol Syst. 2014;45: 371–395. doi:10.1146/annurev-ecolsys-120213-091627

5. Bridge D, Cunningham CW, Schierwater B, DeSalle R, Buss LW. Class-level relationships in the phylum Cnidaria: evidence from mitochondrial genome structure. Proc Natl Acad Sci U S A. 1992;89: 8750–8753. Available:http://www.ncbi.nlm.nih.gov/pubmed/1356268

6. Berntson EA, France SC, Mullineaux LS. Phylogenetic relationships within the class Anthozoa (phylum Cnidaria) based on nuclear 18S rDNA sequences. Mol Phylogenet Evol. 1999;13: 417–433. doi:10.1006/mpev.1999.0649

7. Collins AG, Schuchert P, Marques AC, Jankowski T, Medina M, Schierwater B. Medusozoan phylogeny and character evolution clarified by new large and small subunit rDNA data and an assessment of the utility of phylogenetic mixture models. Syst Biol. 2006;55: 97–115. doi:10.1080/10635150500433615

8. Kayal E, Roure B, Philippe H, Collins AG, Lavrov DV. Cnidarian phylogenetic relationships as revealed by mitogenomics. BMC Evol Biol. 2013;13: 5. doi:10.1186/1471-2148-13-5

9. Marques AC, Collins AG. Cladistic analysis of Medusozoa and cnidarian evolution. Invertebr Biol. Blackwell Publishing Ltd; 2004;123: 23–42. doi:10.1111/j.1744-7410.2004.tb00139.x

10. Bridge D, Cunningham CW, DeSalle R, Buss LW. Class-level relationships in the phylum Cnidaria: molecular and morphological evidence. Mol Biol Evol. 1995;12: 679–689. Available: http://www.ncbi.nlm.nih.gov/pubmed/7659022

11. McFadden CS, France SC, Sánchez JA, Alderslade P. A molecular phylogenetic analysis of the Octocorallia (Cnidaria: Anthozoa) based on mitochondrial protein-coding sequences. Mol Phylogenet Evol. 2006;41: 513–527. doi:10.1016/j.ympev.2006.06.010

12. Cartwright P, Evans NM, Dunn CW, Marques AC, Miglietta MP, Schuchert P, et al. Phylogenetics of Hydroidolina (Cnidaria: Hydrozoa). J Mar Biol Assoc UK. 2008;88: 1663–1672.

13. Rodríguez E, Barbeitos MS, Brugler MR, Crowley LM, Grajales A, Gusmão, L et al. Hidden among sea anemones: the first comprehensive phylogenetic reconstruction of the order Actiniaria (Cnidaria, Anthozoa, Hexacorallia) reveals a novel group of hexacorals. PLoS One. 2014;9: e96998. doi:10.1371/journal.pone.0096998

14. Cartwright P, Halgedahl SL, Hendricks JR, Jarrard RD, Marques AC, Collins AG, et al. Exceptionally preserved jellyfishes from the Middle Cambrian. PLoS One. 2007;2: e1121. doi:10.1371/journal.pone.0001121

15. Cartwright P, Collins A. Fossils and phylogenies: integrating multiple lines of evidence to investigate the origin of early major metazoan lineages. Integr Comp Biol. 2007;47: 744–751. doi:10.1093/icb/icm071

16. Park T-Y, Woo J, Lee D-J, Lee D-C, Lee S-B, Han Z, et al. A stem-group cnidarian described from the mid-Cambrian of China and its significance for cnidarian evolution. Nat Commun. 2011;2: 442. doi:10.1038/ncomms1457

17. Stampar SN, Maronna MM, Kitahara MV, Reimer JD, Morandini AC. Fast-evolving mitochondrial DNA in Ceriantharia: a reflection of hexacorallia paraphyly? PLoS One. 2014;9: e86612. doi:10.1371/journal.pone.0086612

18. Daly M, Brugler MR, Cartwright P, Collins AG, Dawson MN, Fautin DG, et al. The phylum Cnidaria: A review of phylogenetic patterns and diversity 300 years after Linnaeus. Zootaxa. Magnolia Press; 2007;1668: 127–182. Available: https://kuscholarworks.ku.edu/handle/1808/13641

19. France SC, Rosel PE, Agenbroad JE, Mullineaux LS, Kocher TD. DNA sequence variation of mitochondrial large-subunit rRNA provides support for a two-subclass organization of the Anthozoa (Cnidaria). Mol Mar Biol Biotechnol. 1996;5: 15–28. Available: http://www.ncbi.nlm.nih.gov/pubmed/8869515

20. Odorico DM, Miller DJ. Internal and external relationships of the Cnidaria: implications of primary and predicted secondary structure of the 5’-end of the 23S-like rDNA. Proc Biol Sci. 1997;264: 77–82. doi:10.1098/rspb.1997.0011

21. Song J, Won JH. Systematic relationship of the anthozoan orders based on the partial nuclear 18S rDNA sequences. Korean J Biol Sci. 1997;1: 43–52. doi:10.1080/12265071.1997.9647347

22. Won J, Rho B, Song J. A phylogenetic study of the Anthozoa (phylum Cnidaria) based on morphological and molecular characters. Coral Reefs. Springer-Verlag; 2001;20: 39–50. doi:10.1007/s003380000132

23. Reft AJ, Daly M. Morphology, distribution, and evolution of apical structure of nematocysts in hexacorallia. J Morphol. 2012;273: 121–136. doi:10.1002/jmor.11014

24. Bayer FM. Octocorallia. Treatise on invertebrate paleontology. University of Kansas Press Lawrence; 1956; 166–231.

25. Daly M, Fautin DG, Cappola VA. Systematics of the Hexacorallia (Cnidaria: Anthozoa). Zool J Linn Soc. Blackwell Science Ltd; 2003;139: 419–437. doi:10.1046/j.1096-3642.2003.00084.x

26. Park E, Hwang D-S, Lee J-S, Song J-I, Seo T-K, Won Y-J. Estimation of divergence times in cnidarian evolution based on mitochondrial protein-coding genes and the fossil record. Mol Phylogenet Evol. 2012;62: 329–345. doi:10.1016/j.ympev.2011.10.008

27. Beagley CT, Macfarlane JL, Pont-Kingdon GA, Okimoto R, Okada NA, Wolstenholme DR. Mitochondrial genomes of anthozoa (Cnidaria). Prog Cell Cycle Res. 1995;5: 149–153.

28. Bilewitch JP, Degnan SM. A unique horizontal gene transfer event has provided the octocoral mitochondrial genome with an active mismatch repair gene that has potential for an unusual self-contained function. BMC Evol Biol. 2011;11: 228. doi:10.1186/1471-2148-11-228

29. Chen CA, Odorico DM, Tenlohuis M, Veron JEN, Miller DJ. Systematic Relationships within the Anthozoa (Cnidaria: Anthozoa) Using the 5’-end of the 28S rDNA. Mol Phylogenet Evol. 1995;4: 175–183. doi:10.1006/mpev.1995.1017

30. Berntson EA, Bayer FM, McArthur AG, France SC. Phylogenetic relationships within the Octocorallia (Cnidaria: Anthozoa) based on nuclear 18S rRNA sequences. Mar Biol. Springer-Verlag; 2001;138: 235–246. doi:10.1007/s002270000457

31. McFadden CS, Sánchez JA, France SC. Molecular phylogenetic insights into the evolution of Octocorallia: a review. Integr Comp Biol. 2010;50: 389–410. doi:10.1093/icb/icq056

32. Mariscal RN. Nematocysts. Coelenterate biology: reviews and new perspectives. Academic Press: New York, NY, USA; 1974; 129–178.

33. Collins AG. Phylogeny of Medusozoa and the evolution of cnidarian life cycles. J Evol Biol. Blackwell Science Ltd; 2002;15: 418–432. doi:10.1046/j.1420-9101.2002.00403.x

34. Dunn CW. Complex colony-level organization of the deep-sea siphonophore Bargmannia elongata (Cnidaria, Hydrozoa) is directionally asymmetric and arises by the subdivision of pro-buds. Dev Dyn. 2005;234: 835–845. doi:10.1002/dvdy.20483

35. Van Iten H, de Moraes Leme J, Simões MG, Marques AC, Collins AG. Reassessment of the phylogenetic position of conulariids (?Ediacaran-Triassic) within the subphylum medusozoa (phylum cnidaria). J Syst Palaeontol. 2006;4: 109–118. doi:10.1017/S1477201905001793

36. Dunn CW, Howison M, Zapata F. Agalma: an automated phylogenomics workflow. BMC Bioinformatics. 2013;14: 330. doi:10.1186/1471-2105-14-330

37. Stamatakis A, Aberer AJ. Novel Parallelization Schemes for Large-Scale Likelihood-based Phylogenetic Inference. 2013 IEEE 27th International Symposium on Parallel and Distributed Processing. IEEE; pp. 1195–1204. doi:10.1109/IPDPS.2013.70

38. Lanfear R, Calcott B, Ho SYW, Guindon S. Partitionfinder: combined selection of partitioning schemes and substitution models for phylogenetic analyses. Mol Biol Evol. 2012;29: 1695–1701. doi:10.1093/molbev/mss020

39. Lartillot N, Rodrigue N, Stubbs D, Richer J. PhyloBayes MPI. Phylogenetic reconstruction with infinite mixtures of profiles in a parallel environment. Syst Biol. 2013; doi:10.1093/sysbio/syt022

40. Lartillot N, Philippe H. A Bayesian mixture model for across-site heterogeneities in the amino-acid replacement process. Mol Biol Evol. 2004;21: 1095–1109. doi:10.1093/molbev/msh112

41. Swofford DL, Olsen GJ, Waddell PJ, Hillis DM. {Phylogenetic inference}. In: Hillis DM, Moritz C, Mable BK, editors. Molecular systematics (2nd ed). Sinauer Associates, Inc.; 1996. pp. 407–514. Available: http://www.citeulike.org/group/1390/article/768694

42. Church SH, Ryan JF, Dunn CW. Automation and Evaluation of the SOWH Test of Phylogenetic Topologies with SOWHAT. bioRxiv. 2014; doi:10.1101/005264

43. Philippe H, Derelle R, Lopez P, Pick K, Borchiellini C, Boury-Esnault N, et al. Phylogenomics revives traditional views on deep animal relationships. Curr Biol. 2009;19: 706–712. doi:10.1016/j.cub.2009.02.052

44. Yoffe C, Lotan T, Benayhau Y. A modified view on octocorals: Heteroxenia fuscescens nematocysts are diverse, featuring both an ancestral and a novel type. PLoS One. 2012;7: e31902. doi:10.1371/journal.pone.0031902

45. Werner B. New investigations on systematics and evolution of the class Scyphozoa and the phylum Cnidaria. Publ Seto Mar Biol Lab. 1973;20: 35–61. Available: http://repository.kulib.kyoto-u.ac.jp/dspace/handle/2433/175791

46. Cartwright P, Nawrocki AM. Character evolution in Hydrozoa (phylum Cnidaria). Integr Comp Biol. 2010;50: 456–472. doi:10.1093/icb/icq089

47. Nawrocki AM, Cartwright P. A novel mode of colony formation in a hydrozoan through fusion of sexually generated individuals. Curr Biol. 2012;22: 825–829. doi:10.1016/j.cub.2012.03.026

48. Dunn CW, Pugh PR, Haddock SHD. Molecular phylogenetics of the siphonophora (Cnidaria), with implications for the evolution of functional specialization. Syst Biol. 2005;54: 916–935. doi:10.1080/10635150500354837

49. Nawrocki AM, Collins AG, Hirano YM, Schuchert P, Cartwright P. Phylogenetic placement of Hydra and relationships within Aplanulata (Cnidaria: Hydrozoa). Mol Phylogenet Evol. 2013;67: 60–71. doi:10.1016/j.ympev.2012.12.016

